# Which senses do wild vervet monkeys (*Chlorocebus pygerythrus*) use for evaluating potential food items?

**DOI:** 10.64898/2026.03.27.714682

**Authors:** Clara Ondina Ferreira da Silva Teixeira, Erica van de Waal, Matthias Laska, Alba Motes Rodrigo

**Author notes:** Corresponding author Alba Motes-Rodrigo, Department of Ecology and Evolution, University of Lausanne, Lausanne, Switzerland.

## Abstract

Traditionally, primates have been considered primarily visual animals. However, studies across a variety of taxa suggest that, in the context of food evaluation, the reliance on this sense might be more nuanced that previously thought, with dietary specialization and food item properties leading to differences in sensory prioritization. We performed a field-based study assessing the use of sensory cues during food evaluation as well as food-related behaviours such as muzzle contact in two mixed-sex groups of wild vervet monkeys including three age classes over a period of five months (n_monkeys_ = 44). Using a total of 18868 food evaluation observations collected over 44 hours of focal follows, we found that vervets mainly relied on their sense of vision when evaluating food (96.8% of all instances). Sensory usage varied according to food category and sex differences were only observed in the use of smell for a subset of these. Juveniles initiated muzzle contact and used tactile inspection more often than adults whereas females received muzzle contact more often than males. In addition, the low rejection rates suggest that most food items were familiar to the vervets regardless of age and sex. These findings are in line with optimal foraging theory according to which the food evaluation process should be adapted to the familiarity of food items and allows individuals to maximize their intake of energy and critical nutrients, while minimizing the time and effort in food evaluation.

## Introduction

Primates have traditionally been considered as primarily visual animals (Kaas and Balaram, 2014; Martinez-Trujillo and Piza, 2026). However, it seems reasonable to assume that primates do not only rely on visual information but also use other senses in a variety of behavioral contexts, including the evaluation of potential food items. Indeed, non-visual stimuli such as smell, taste and touch have repeatedly been reported to play a role in primate food selection (Veilleux et al., 2022). This should not be surprising considering that fruits consumed by primates, for example, systematically change in their odor (Nevo et al., 2022) and taste composition (Rodrigo et al., 2012) as well as in their hardness (Kinzey and Norconk, 1990) during maturation and thus provide honest signals of fruit ripeness and in turn, of nutritional value (Nevo et al., 2015). Primates have also been reported to use olfactory cues to recognize whether a potential food item is putrefying or rotting and thus to make informed decisions about its consumption or rejection (Laska et al., 2007a; Laska and Hernandez-Salazar, 2015).

So far, only a limited number of studies have directly assessed which sensory cues free-ranging nonhuman primates use to inspect and select food items and whether their reliance on the different senses depends on food familiarity and/or dietary specialization. Laska et al. (2007b) reported that captive squirrel monkeys (*Saimiri sciureus*) and spider monkeys (*Ateles geoffroyi*) used olfactory, gustatory, and tactile cues in addition to visual information to evaluate novel foods. However, as soon as these novel foods became increasingly familiar due to repeated presentation, the animals relied primarily on visual inspection prior to consumption, suggesting that familiarity may play an important role in the use of different senses for food selection in primates. Additionally, the two species were found to differ in their relative use of non-visual cues. Spider monkeys relied more on olfactory cues when evaluating novel foods than squirrel monkeys, which relied more heavily on tactile cues. Similarly, Sanchez-Solano et al. (2022) found that free-ranging mantled howler monkeys (*Alouatta palliata*) used non-visual sensory cues in food selection, particularly when assessing fruits which do not change color during ripening. In this species, the relative use of smell and touch information, respectively, changed as a function of ripeness of the fruits. Melin et al. (2022) reported that free-ranging frugivorous spider monkeys used olfactory cues most often among the non-visual senses when assessing potential food items whereas omnivorous capuchin monkeys (*Cebus imitator*) used tactile cues most often. This suggests that dietary specialization may affect the use of the different senses for food selection in primates.

Vervet monkeys (*Chlorocebus pygerythrus*) have been described as “opportunistic omnivores” and consume a wide variety of food types such as fruits, leaves, flowers, seeds, and animal matter including arthropods, birds and bird eggs, and even small mammals into their diet (Struhsaker, 1967; Tournier et al., 2014; Brun et al., 2022). Despite this detailed knowledge about the species’ diet composition, little is known about which senses vervet monkeys use for food selection nor whether their use varies according to age and sex. Previous behavioral studies found that vervet monkeys perform different food-opening and food-cleaning techniques (van de Waal et al., 2012; Canteloup et al., 2021) which infant vervets acquire from their mothers (van de Waal et al., 2014). These findings imply elaborate manipulation of food items prior to consumption which could lead to frequent use of tactile cues by younger individuals during food selection. Vervet monkeys have also been reported to engage in muzzle contact where one individual brings its nose and mouth area into close proximity to that of another individual (Nord et al., 2021; Dongre et al., 2024). This behavior, which is often initiated by infants and juveniles and directed towards adults, is thought to serve as a means of obtaining smell and taste information about the food item that the adult is consuming or has recently consumed (Laidre 2009). In support of this potential information-gathering hypothesis, Dongre et al. (2024) found that during novel food experiments, muzzle contact was directed from uninformed to knowledgeable individuals. Therefore, both food cleaning and muzzle contact behaviors suggest that vervet monkeys may employ non-visual sensory cues for food selection.

Given the paucity of data on which senses primates from different age and sex classes use for evaluating potential food items, the aims of the present study were to 1) record and analyze the use of visual, tactile, olfactory, and gustatory cues during the evaluation of potential food items in free-ranging vervet monkeys, 2) assess whether the use of the different senses differs as a function of the type of food, 3) investigate whether sex and age influence the use of different senses during food evaluation and 4) assess whether food-related behaviours (food preparation, muzzle contact and food rejection) vary as a function of age and sex.

## Methods

### Animals

The study took place at the iNkawu Vervet Project, a long-term field site for the study of vervet monkeys (*Chlorocebus pygerythrus*) located in a private reserve called Mawana Game Reserve in KwaZulu-Natal, South Africa. The field site hosts seven free-ranging groups of vervet monkeys habituated to the presence of humans. We observed all individuals from two groups,, Baie Dankie (BD) and Ankhase (AK). Our sample included 20 females (12 from BD and 8 from AK) and 24 males (16 from BD and 8 from AK), with 3 animals being infants (<1 year of age), 15 being juveniles (for males: 1 year > dispersal; for females: < 3 years old), and 26 being adults (for males: post-dispersal; for females: >3 years old).

### Data collection

Over a period of five months (August-December 2024) during the transition between the dry and the wet seasons, we observed each vervet monkeys in the two target groups using focal animal sampling (Altmann, 1974). We recorded each instance of a focal animal inspecting a potential food item and the occurrence of visual, tactile, olfactory, and gustatory evaluation (see ethogram, Table 1). Further, we recorded the type of potential food item (fruits, leaves, flowers, seeds, bark, gum, roots, arthropods, and unknown) that an animal inspected, and whether the inspected potential food item was consumed or rejected by an animal following its evaluation. Finally, we recorded all instances of mouth-to-mouth/muzzle contacts between dyads of individuals while one of them was feeding on a food item. Each of the 44 individuals was observed for a total of 60 minutes of feeding behavior, spread across the whole 5-month period.

**Table 1.** Ethogram of the behaviors considered in this study.

| **Behaviors** | **Description** |
| --- | --- |
| Vision | An animal focusing its field of vision on a potential food item for at least 3 seconds |
| Olfaction | An animal bringing a food item close to (<5 cm) or in contact with its nose or performing nose movements indicative of sniffing |
| Taste | An animal licking a food item or taking a first bite as a probe |
| Touch | An animal taking a food item into its hands and manipulating it or biting on it with its teeth without removing a part of the food item |
| Food preparation | An animal scrubbing a food item on a substrate or using its hands to clean it or peeling a food item using its hands and/or teeth |
| Food consumption | An animal putting a food item into its mouth and swallowing it with or without chewing |
| Food rejection | An animal discarding a food item after its inspection or after a first probing bite |
| Performing muzzle contact | An animal bringing its muzzle into close proximity (<5 cm) to that of a conspecific consuming or having consumed a food item |
| Receiving muzzle contact | Another animal bringing its muzzle into close proximity (<5 cm) to that of a conspecific consuming or having consumed a food item |

### Ethical note

The experiments reported here comply with the *American Society of Primatologists’ Principles for the Ethical Treatment of Primates*, with the *European Union Directive on the Protection of Animals Used for Scientific Purposes* (EU Directive 2010/63/EU), with the ARRIVE guidelines, and with current Swedish, Swiss, and South African animal welfare laws. Given the non-invasive, observational nature of the study, no specific ethical approval beyond the consent of the field-site director and the landowners was required to perform data collection.

### Data analysis

All data analysis was performed with R and RStudio (R Core Team, 2021). The Shapiro-Wilk test showed that our data were not normally distributed. To assess possible differences in the frequency of use between the four senses we used Friedman ANOVA. When the ANOVA yielded a significant outcome, we conducted a post-hoc analysis using the Wilcoxon signed-rank test to determine which specific pairs of senses differed significantly from each other.

To examine how sensory modality use varied across food categories and between sexes and age classes, we fitted generalised linear mixed models (GLMMs) with group as control predictor and monkey identity as a random intercept effect to account for repeated measures. The response variable was the raw count of observations of each sense used per individual per food category. We first assessed which model complexity was supported by the data for each sense by examining whether at least two combinations of sex and age class had non-zero observations for each food category (vision = 7, smell = 4, taste = 4, touch = 5). All models were initially fitted with a Poisson error distribution using the lme4 package in R. Residuals were inspected for overdispersion and zero-inflation using the DHARMa package. Post-hoc pairwise comparisons were conducted using the emmeans package with Bonferroni correction. Proportions shown in figures were calculated as the number of observations of a given sense while eating a given food divided by the total sensory observations recorded for that individual eating that food.

To assess individual variation in food-related behaviours, we modelled rejection, preparation, muzzle contact initiation, and muzzled contact reception as a function of sex and age class. Muzzle contact initiation and reception were modelled as binary outcomes (present/absent) using binomial GLMs due to data sparsity. Rejection and preparation counts were initially modelled using Poisson GLMs, but both showed significant overdispersion and zero-inflation (DHARMa tests, all p < 0.001) and thus were refit as zero-inflated negative binomial models using package glmmTMB.

All data and code used to perform the analyses and create the figures can be found in https://github.com/AlbaMotes/vervet_food_inspection_senses.git.

## Results

### Use of the senses when evaluating potential food items

We recorded a total of 18868 instances of the use of the senses when a vervet monkey evaluated a potential food item. The use of the sense of vision accounted for 96.8% (18258 instances) of all observations, touch for 1.8% (334), olfaction for 1.1% (203), and taste for 0.4% (73), respectively (Figure 1). Accordingly, vision was used significantly more often by the vervet monkeys than the other three senses (χ^2^-test, p < 0.001 with all three comparisons), touch was significantly used more often than olfaction and taste (χ^2^-test, p < 0.01 with both comparisons), and olfaction was used significantly more often than taste (χ^2^-test, p < 0.01).

**Figure 1:**
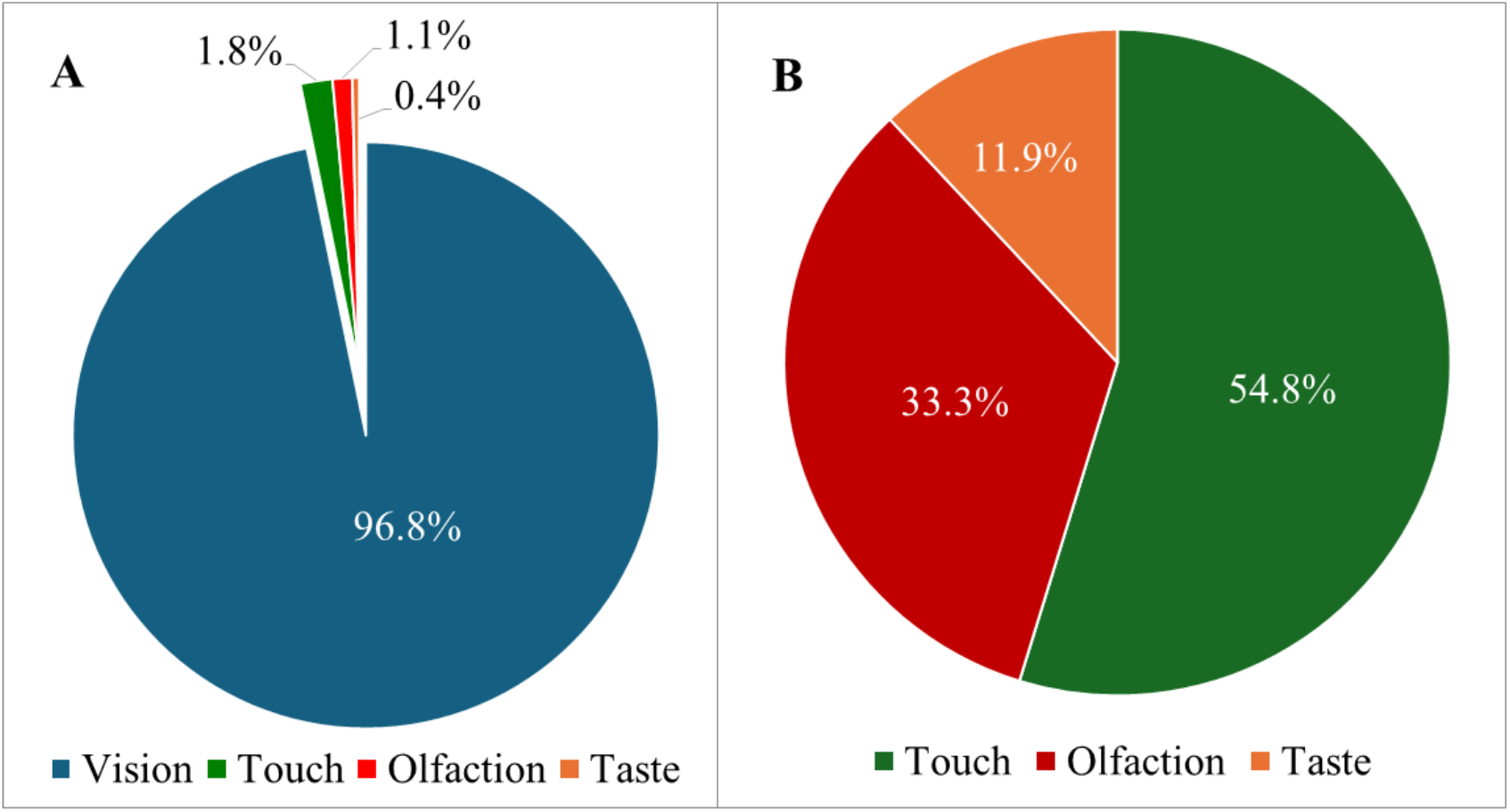
Distribution of the proportions at which the vervet monkeys (n = 44) used the different senses during food evaluation. Panel A: frequency distribution of all four senses. Panel B: frequency distribution of the non-visual senses.

### Sex and Age differences in the use of the senses to evaluate different food types

Leaves were the most frequently inspected category of recognizable food and accounted for 38.1% (7326 instances) of all observations when a vervet monkey inspected a potential food item. Bark accounted for 9.9% of all observations (1899 instances), fruits for 7.5% (1443 instances), flowers for 4.5% (865 instances), arthropods for 1.6% (311 instances), gum for 0.2% (43 instances), roots for 0.08% (16 instances), seeds for 0.04% (7 instances), and fungi for 0.02% (4 instances) of all observations, respectively. In 38.0% of all observations (7318 instances) we failed to identify the category of the inspected food item which therefore was recorded as unknown.

No overall sex or age differences were found in the use of vision (vision_sex: χ^2^ = 1.36, p = 0.24; vision_age: χ^2^ = 0.95, p = 0.62). However, the use of this sense varied according to the food item consumed (χ^2^ = 442.59, p < 0.001; Figure 2). While the differences among food categories were small in absolute terms and close to ceiling, leaves and unknown foods were visually inspected significantly more often than all other food categories, while flowers and gum were the least inspected (Figure 2).

**Figure 2.**
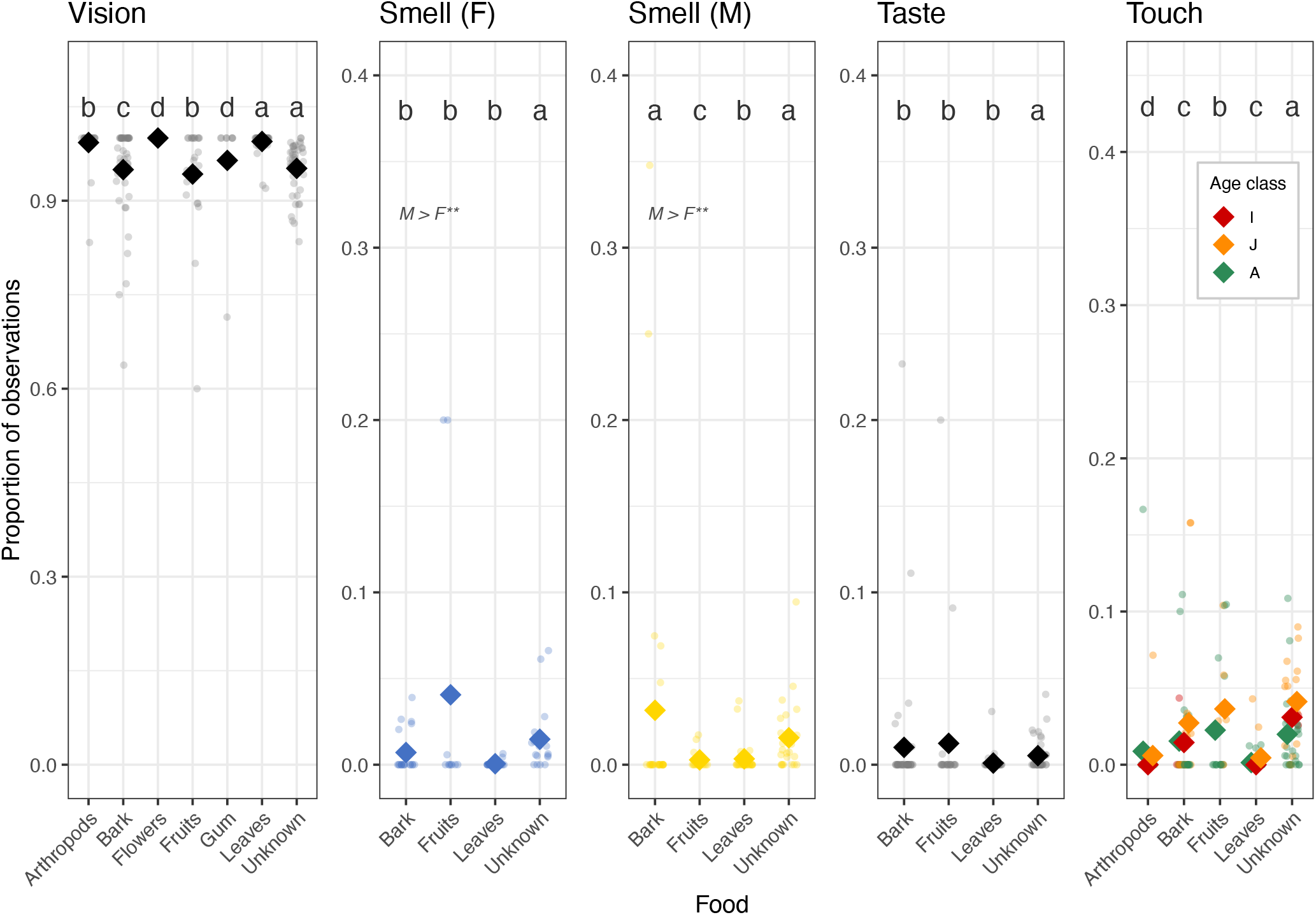
Proportion of observations of sense usage for a specific food category per individual. The data on use of smell for food evaluation is split by sex reflecting the significant interaction between sex and food type. Females (F) are represented in blue, and males (M) are represented in yellow. Diamonds represent mean proportion of sense usage per food category. In the touch panel, colors represent the different age classes and diamonds represent the age class mean (I = infant, J = juvenile, A = adult). Asterisks denote significant differences (**p< 0.005) and letters indicate the results of pairwise comparisons among food categories with Bonferroni corrected p-values.

The use of smell was influenced by both food item consumed and sex in interaction (χ^2^ = 22.70, p < 0.001), meaning that how much smell was used per sex differed depending on the food item consumed. Males and females differed significantly in smell use when inspecting bark, with males using smell more than females (estimate = -1.52, SE = 0.55, z = -2.79, p = 0.005). Among females, smell was used significantly more when inspecting unknown food compared to bark, fruits, and leaves (all p < 0.001). Among males, smell was mostly used for inspecting bark and unknown foods, over leaves (p < 0.001) and fruits (p < 0.001). Age classes did not differ in the use of smell (χ^2^ = 2.12, p = 0.35).

The use of taste varied significantly by food item (χ^2^ = 28.55, p < 0.001), but pairwise comparisons revealed very small differences among food types, which means were very close to 0 (Figure 2). No overall sex or age differences were found in the use of taste (taste_sex: χ^2^ = 0.47, p = 0.49; taste_age: χ^2^ = 1.83, p = 0.40).

Touch use varied significantly by food item (χ^2^ = 133.19, p < 0.001) and age class (χ^2^ = 11.59, p = 0.003). Unknown food items were explored via touch more than any other food category (all p < 0.001), and juveniles used touch significantly more than adults (ratio = 0.47, SE = 0.10, z = -3.40, p = 0.002). No overall sex differences were found in the use of touch (χ^2^ = 0.06, p = 0.81).

### Consumption and rejection of food items

After inspection, potential food items were consumed in 98.9% of all observations (17120 instances) and rejected in only 1.1% of all observations (186 instances). Males and females did not differ significantly from each other in their rejection rates (χ^2^ = 0.002, p = 0.96). Similarly, the three age classes did not differ significantly from each other in their rejection rates (χ^2^ = 4.82, df = 2, p = 0.09). Among the different food categories, fruits were rejected most often (2.4% of cases), followed by unknown (1.7%), bark (1.6%) and leaves (0.3%). All other food categories (flowers, seeds, gum, root, and arthropods) were never rejected.

### Preparation of food items

We observed 222 instances of food preparation, defined as an animal scrubbing a food item on a substrate or using its hands to clean it or peeling a food item using its hands and/or teeth. This corresponds to 0.6% of all observations of a vervet monkey inspecting a potential food item. Among the different food categories, roots were prepared most often (3.1% of cases), followed by arthropods (1.8%), unknown (1.3%), bark (0.3%), fruit (0.3%) and leaves (0.2%). The remaining food categories (flowers, seeds, and gum) were never prepared. Males and females did not significantly differ in their food preparation rates (χ^2^ = 0.19, p = 0.66). Similarly, the three age classes did not significantly differ in their food preparation rates (χ^2^ = 2.00, df = 2, p = 0.37).

### Muzzle contact

We observed a total of 54 instances of muzzle contact, with 30 cases in which the focal animal was the recipient, and 24 cases in which the focal animal was the initiator of muzzle contact. Age class significantly influenced the probability of initiating muzzle contact (χ^2^ = 6.68, p = 0.035), with adults showing a trend towards initiating less contacts than infants and juveniles, though posthoc pairwise comparisons did not reach significance after Bonferroni correction (all p > 0.11). Sex did not significantly influence initiation (χ^2^ = 0.99, p = 0.32). Regarding reception, females were significantly more likely to receive muzzle contact than males (OR = 4.34, SE = 2.89, p = 0.027), while age class had no effect (χ^2^ = 0.51, p = 0.77).

## Discussion

The results of the present study demonstrate that free-ranging vervet monkeys mainly rely on their sense of vision when evaluating potential food items. Among the non-visual senses, they relied most often on touch, followed by smell and taste and their frequency varied as a function of the type of food. In addition, we found sex differences in the use of smell for bark inspection as well as in the frequency at which individuals received muzzle contact. Juveniles used touch more often than adults and, together with infants, they were the most likely to initiate muzzle contact.

### Use of the senses when evaluating potential food items

Our findings that the vervet monkeys of the present study mainly relied on their sense of vision when evaluating potential food items should be expected considering their high visual acuity and trichromatic vision (Martinez-Trujillo and Piza, 2026). Further, vision does not only provide information about the chromatic (i.e., hue and saturation) and luminance (i.e., brightness) properties of potential food items but also about their shape, size, pattern, and movement (Veilleux et al., 2022). Accordingly, this variety of color- and non-color-related visual cues may be sufficient for vervet monkeys to make informed decisions about the palatability and quality of familiar food items in their environment whereas non-visual cues may be more prominently explored with unfamiliar food. Studies on other primate species support this notion as squirrel monkeys and spider monkeys, for example, have been reported to use olfactory, gustatory, and tactile cues when evaluating novel foods, but almost exclusively used visual cues when foods were familiar (Laska et al., 2007a). Similarly, orangutans, chimpanzees, bonobos and tufted capuchins were all found to use non-visual cues more often when presented with novel foods compared to familiar foods (Forss et al., 2019; Visalberghi et al., 2003). Adapting the food evaluation process according to the familiarity with a given food item is in line with optimal foraging theory as natural selection should favor individuals that succeed in maximizing their intake of energy and critical nutrients, and this should be reflected in minimizing the time and effort that an animal invests into food evaluation (Stephens et al., 2008).

With regard to the non-visual senses, we found that vervet monkeys used their sense of touch significantly more often than their chemical senses, and among the latter they used their sense of smell significantly more often than their sense of taste when evaluating potential food items. Several hypotheses have been put forward trying to explain between-species differences in the relative use of the non-visual senses in primates in the context of food selection. Some authors argue that the degree of manual dexterity may be linked to a species’ use of the sense of touch (Heldstab et al., 2016). Platyrrhine primates, with the exception of capuchins, lack independent motor control of their fingers which prevents them from performing a precision grip and thus limits their ability of fine food manipulation with their hands (Verendeev et al., 2016). Catarrhine primates such as vervet monkeys, in contrast, possess this motor ability and are thus able to perform fine manipulations of food items. Our observation that the vervet monkeys of the present study repeatedly displayed manual food preparation by scrubbing or cleaning or peeling a food item prior to consumption fits to this notion. A previous study reported that they even perform different food-cleaning techniques (van de Waal et al., 2012), lending further support to the presumed importance of the sense of touch for the evaluation of food.

Other authors argue that the relative size of the olfactory bulbs may reflect a species’ reliance upon its sense of smell (Barton, 2006). Platyrrhine primates are known to have larger olfactory bulbs relative to total brain size compared to catarrhine primates (Heritage, 2014). This supports our finding that the vervet monkeys of the present study used their sense of smell only rarely when evaluating food and significantly less often than their sense of touch. Nevertheless, vervet monkeys have been reported to perform scent marking as a means of social communication (Freeman et al., 2012) and to smell at the genitals of female conspecifics to assess their reproductive status (Brain, 1965). This suggests that they use olfactory cues in different behavioral contexts and thus that their sense of smell should be sufficiently acute to also serve a function in the evaluation of food.

Previous studies have proposed that dietary specialization may affect the use of the non-visual senses in primate food selection (Dominy et al., 2001). Indeed, some studies found that frugivorous species such as spider monkeys used their sense of smell more often when evaluating food items than folivorous species such as howler monkeys or omnivorous species such as capuchins (Laska et al., 2007a; Sanchez-Solano et al., 2022; Melin et al., 2022). This seems plausible as fruits consumed by primates are known to change their odor during maturation and thus provide information about their ripeness and nutritional value whereas leaves usually do not (Nevo et al., 2015; Nevo et al., 2022). Vervet monkeys are considered as “opportunistic omnivores” feeding on a broad diet which includes only a relatively small proportion of fruits (Tournier et al., 2014; Brun et al., 2022). Our finding that less than 10% of all food items inspected by the vervet monkeys of the present study were fruits fits to this notion and may also explain why they used olfactory cues only rarely when inspecting food items.

Finally, our finding that the vervet monkeys used their sense of taste significantly less often than their senses of touch and smell when evaluating potential food items may be explained by the notion that gustation can be considered as the “final sentinel” against the ingestion of unpalatable or toxic food (Glendinning, 2021). Considering that food avoidance learning in primates is extraordinarily quick and robust (Laska and Metzker, 1998), and thus that a novel food may become familiar to an animal after only one episode of ingestion, the warning function of the sense of taste may be restricted to the evaluation of novel food items. This notion is in line with our finding that the vervet monkeys of the present study rejected food items after inspection in only 1.1% of all cases, suggesting that the vast majority of the food items that they consumed were familiar to them, rendering the use of taste cues for food evaluation unnecessary.

### Sex differences in the evaluation of potential food items

We found limited evidence for differences in sensory use between male and female vervet monkeys. Whereas sex differences in diet composition (e.g., Koch et al., 2017) and foraging strategies (e.g., Bean, 2004) have been described in various primate species which can be plausibly explained by differences in energy and nutrient demand associated with reproduction and infant care, there is no *a priori* reason to assume that males and females should differ in their overall use of the senses when evaluating food. The only statistically significant difference between the sexes that we found was in their use of smell when evaluating bark. At this point we can only speculate whether this difference might be attributed to possible differences between males and females in their need to avoid the ingestion of too high levels of plant secondary compounds or to meet their need for certain macro- or micronutrients which may be particularly abundant in this type of food. While the interpretation of these results would benefit from further investigation, our findings are the first to suggest potential context-dependent differences in sensory evaluation between the sexes.

We also found that males and females did not differ significantly from each other in their rates of food rejection. A marked increase in food rejection rates has been reported to occur in female primates during pregnancy (Czaja, 1975). The fact that only few of the female vervet monkeys of the present study were visibly pregnant or gave birth during our observation period might explain that their food rejection rates resembled those of their male conspecifics.

Finally, we also found no significant differences in the rates at which male and female vervet monkeys prepared their food by scrubbing, cleaning or peeling a food item prior to consumption. This finding is in line with a previous study which reported that male and female vervet monkeys did not differ in the frequency and type of their food cleaning techniques (van de Waal et al., 2012).

### Age differences in the evaluation of potential food items

We found that age classes only differed in the use of touch, with juveniles using this sense more often than adults. These results could be explained by differences in pre-existing dietary knowledge among individuals that would lead juveniles and infants to rely more heavily than adults on their non-visual senses (Fragaszy et al., 1997; Nord et al., 2022). The lack of age effects in the use of the other sense could be related to the high degree of familiarity of the food items consumed. Further studies on sensory use that involve presenting novel foods or modified familiar foods to vervet monkeys could test this hypothesis.

We found that infant, juvenile, and adult vervet monkeys did not significantly differ from each other in their food preparation rates. In this context, it is interesting to note that infant vervet monkeys have been shown to have the capacity for detailed copying and thus for early learning of food-cleaning techniques from their mothers and matriline members (van de Waal et al., 2014).

Given that approximately 30% of the plant species present in the home ranges of the studied groups are toxic (van de Waal pers. comm.), we expected higher rejection rates. The fact that rejection rates were below 5% suggest that vervets avoid these plants altogether, indicating local knowledge on food toxicity. Our negative results regarding age differences in food rejection rates may be surprising, as infants might be expected to be cautious or even neophobic towards novel objects, situations, and foods due to their vulnerability and lack of experience (Janson and van Schaik, 1993). However, infant vervet monkeys have also been reported to display shorter latencies to approach novel objects and foods than adults, are generally more willing to take risks than older conspecifics (Fairbanks 1993) and have high innovation rates (Dongre et al., 2024). Further, in contrast to juveniles and adults, infant vervet monkeys have been found to lack distinct preferences for certain food categories and instead treated all types of food as equally appealing (Hauser, 1993).

### Muzzle contact

Muzzle contact involves one individual bringing its nose and mouth area into close proximity to that of a conspecific (Laidre, 2009). Previous studies have shown that this behavior is often initiated by infants or juveniles and directed towards adults as is thought to serve as a means of obtaining smell information about the food item that the adult is consuming or has recently consumed (Nord et al., 2021). Accordingly, this behavior can be considered as a form of olfactory social learning (Lycett and Henzi, 1992). We found a significant age effect, where infant and juvenile vervet monkeys indeed initiated muzzle contact almost three times more often than adults. We also found that females received muzzle contact significantly more often than males. This could be explained by females being the philopatric sex and therefore having local knowledge on food edibility, which would be in agreement with previous studies that found females to be the preferred role models under experimental conditions (van de Waal et al., 2010). Similarly, an earlier study on vervet monkeys also reported that muzzle contacts were initiated most often by naïve individuals and were targeted most often towards knowledgeable conspecifics (Dongre et al., 2024). The same study also found that muzzle contacts decreased in frequency across repeated exposure to novel foods, supporting the notion of a fast and robust olfactory social learning process. It is interesting to note that, except for vervet monkeys, muzzle contact has so far only been reported in a few other catarrhine primate species including mandrills (*Mandrillus sphinx*), drills (*Mandrillus leucophaeus*), olive baboons (*Papio anubis*), yellow baboons (*Papio cynocephalus*), and Tonkean macaques (*Macaca tonkeana*) (Drapier et al., 2002; Laidre, 2009). All of these species are known to feed on a broad diet and can be described as “opportunistic omnivores”. Future comparative studies should therefore assess whether this type of dietary specialization might be a driving force for the occurrence of muzzle contact as a means of olfactory social learning.

In conclusion, our findings are in line with optimal foraging theory as adapting the food evaluation process according to the familiarity with a given food item should be reflected in minimizing the time and effort that an animal invests into the evaluation of food.

## Acknowledgements

We are grateful to the van der Walt family for giving us the permission to conduct the study on their land at Mawana Game Resrve. We thank the iNkawu Vervet Project for permission to conduct this study and for providing logistic support at the Mawana Private Game Reserve. We are grateful to the field managers, volunteers, and field assistants for their support during data collection and fieldwork. This project was funded by the Swiss National Science Foundation (CRSII-222818) along with the European Research Council under the European Union’s Horizon 2020 research and innovation programme for the ERC ‘KNOWLEDGE MOVES’ starting grant (grant agreement no. 949379) that also supported E.v.d.W.

